# LRIT3 expression in cone photoreceptors restores post-synaptic bipolar cell signalplex assembly and function in *Lrit3^-/-^* mice

**DOI:** 10.1101/2022.08.06.503042

**Authors:** Ronald G. Gregg, Nazarul Hasan, Bart G. Borghuis

**Affiliations:** Department of Biochemistry and Molecular Genetics; Department of Ophthalmology and Visual Sciences; Department of Anatomical Sciences and Neurobiology University of Louisville, Louisville, KY 40292

**Keywords:** Lrit3, congenital stationary night blindness, cone photoreceptors, retina, bipolar cells, retinal ganglion cells, trans-synaptic interactions, LRR proteins, TRPM1

## Abstract

Complete congenital stationary night blindness (cCSNB) is a heterogeneous disorder characterized by poor dim light vision, myopia, and nystagmus, that is caused by mutations in genes critical for signal transmission between photoreceptors and depolarizing bipolar cells (DBCs). One such gene, *LRIT3*, is required for assembly of the post-synaptic signaling complex (signalplex) at the dendritic tips of DBCs, although the number of signalplex components impacted is greater in cone DBCs than in rod bipolar cells. This difference raises the possibility that LRIT3 is expressed both pre- and post-synaptically at cone terminals. Here we show that rAAV-mediated expression of LRIT3 in cones results in robust rescue of cone DBC signalplex components and restores downstream visual function, as measured by the light adapted ERG b-wave and electrophysiological recordings of BCs and RGCs. These data show that LRIT3 successfully restores function to cone DBCs in a trans-synaptic manner, potentially paving the way for therapeutic intervention in LRIT3-associated cCSNB.

## INTRODUCTION

Synaptic function requires the correct alignment and assembly of pre- and postsynaptic proteins. Developing therapies to repair neuronal circuits in disease will benefit from understanding the basic processes that govern synapse assembly. Due to a long history of research on neural circuitry, morphology and, more recently, various Omics, including connectomics, the retina is now at the forefront of efforts aimed at resolving synaptic complex formation and the molecular events required to create functional circuitry.

Vision is initiated when a rod or cone photoreceptor detects a change in luminance and signals this change to postsynaptic cells. Rods mediate vision under dim light conditions and cones under bright light conditions, together encompassing a luminance range that spans about ten log units of intensity. Congenital stationary night blindness (CSNB) represents a heterogeneous group of vision defects that despite no retinal degeneration, result in poor dim light vision, strabismus and myopia (Miyake et al., 1986). There are two main types: Riggs, which results from defects in rod photoreceptor signaling, and Schubert-Bornschein, which can be divided based on the electroretinogram (ERG) into the incomplete (iCSNB) and complete (cCSNB) forms, which are caused by defects in pre-versus post-synaptic mechanisms, respectively(for review see (Zeitz et al., 2015)).

cCSNB is generally caused by defects in the depolarizing bipolar cell (DBC; rod bipolar and ON cone bipolar cell) signalplex, which uses the metabotropic glutamate receptor 6 (mGluR6) (Nakajima et al., 1993) to detect glutamate released from photoreceptors to modulate DBC membrane potential through the TRPM1 channel (Koike et al., 2010; Morgans et al., 2009; Shen et al., 2009). cCSNB results in the absence of both the rod and cone mediated ERG b-waves, because the signalplex in rod and cone DBCs is not functional due to the absence of any one of a number of signalplex members, including mGluR6, GPR179, nyctalopin, TRPM1, or LRIT3 (see review (Furukawa et al., 2020)).

The known composition of the post-synaptic signalplex in the rod BC and cone BCs is similar, however, the identification of LRIT3 as a component of the signalplex assembly revealed a more nuanced picture, showing that requirements for assembly of rod and cone signalplexes differ (Hasan et al., 2018; Hasan et al., 2020; Neuille et al., 2015; Ray, 2013). Rod BCs in LRIT3 deficient mice lose expression of TRPM1 and nyctalopin while the remaining members, mGluR6, GPR179, RGS7/11, and R9AP are expressed and localized to the DBC dendritic tips. In contrast, ON cone BCs in LRIT3 deficient retinas lose all known signalplex proteins from the tips of the dendrites. These differences indicate a more complex interaction in cone versus rod BCs dendrites. Because these changes are all in the post-synaptic DBC signalplex, it was assumed that LRIT3 expression was required in DBCs. We challenged this idea by our demonstration that expression of LRIT3 in rod photoreceptors through transduction with rAAV and a rod-selective rhodopsin gene promoter restores expression of nyctalopin and TRPM1 to rod BCs and recovers their function as measured by the ERG and recordings from BCs and RGCs (Hasan et al., 2019). This unexpected result suggests that LRIT3 can act as a trans-synaptic organizer of the post-synaptic signalplex. Two other studies, one in mouse and one in dog LRIT3 deficient models, restored rod BC signalplex function using postsynaptic targeting of LRIT3 (Miyadera et al., 2022; Varin et al., 2021).

To further explore a trans-synaptic role for LRIT3 in restoring visual function in cCSNB we focused on photopic (cone-mediated) signaling and expressed LRIT3 in cone photoreceptors using sub-retinal injections of rAAV expression vectors with the putative cone specific GNAT2 gene promoter in adult *Lrit3*^*-/-*^ mice. We found that expression of LRIT3 in cone photoreceptors restored expression of the mGluR6/TRPM1/Nyctalopin complex in the dendritic tips of ON cone BCs, and retinal visual function at photopic light levels as measured by the electroretinogram and electrophysiological recordings of ON RGCs. These data show that in addition to its previously established role in organizing the rod to rod bipolar cell synapse, LRIT3 acts also as a trans-synaptic organizer of the signalplex at cone to cone bipolar cell synapses.

## RESULTS

The loss of LRIT3 cause more members of the DBC signalplex in BCs connected to cones than those connected to rods (Hasan *et al*., 2020; Neuille et al., 2017; Ray, 2013). One possibility for this discrepancy is that cones and cone DBCs may both express LRIT3, and pre- and postsynaptic expression is necessary for complete signalplex assembly. To investigate this we expressed LRIT3 in cones using rAAV8 and the ProA1 promoter – a part of the *Gnat2* promoter that was reported to limit expression to cone photoreceptors (Drinnenberg et al., 2018). A schematic of the rAAV construct is shown in Fig.1A and the experimental paradigm in Fig. 1B.

**Figure 1.**
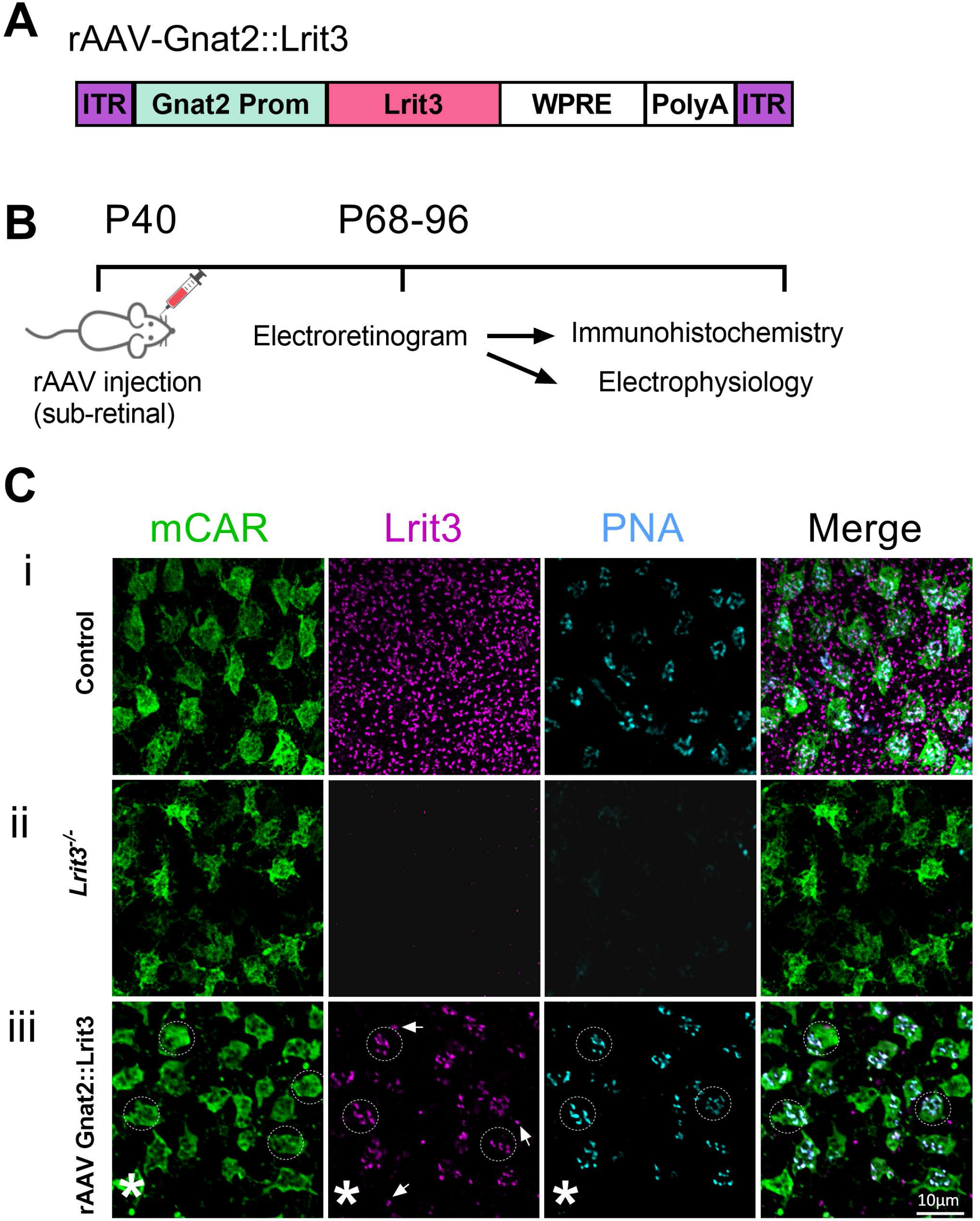
rAAV Gnat2::Lrit3 restores LRIT3 expression to cones. (A) Schematic diagram of the rAAV Gnat2::Lrit3 vector and (B) the time course of treatment and analyses. (C) Retinal whole mounts stained for cone arrestin (mCAR), LRIT3 and PNA imaged at the level of the OPL in (i) control, (ii) *Lrit3*^*-/-*^ and (iii) rAAV Gnat2::Lrit3 treated *Lrit3*^*-/-*^ mouse retinas. Arrowheads in (iii) indicate puncta representing rod terminals. OPL, outer plexiform layer. Scale bar = 10µm. The dashed circles highlight 3 of 22 cone terminals that show restored expression of LRIT3 and PNA. The asterisk highlights a cone that lacks expression of LRIT3 and thus PNA. The infection rate in this region was 17/22 cones (73%).

### rAAV-mediated expression of LRIT3 in *Lrit3*^*-/-*^ cones restores expression of cone DBC signalplex components

Cone DBCs in *Lrit3*^*-/*-^ mice lack all components of the post-synaptic DBC signalplex and cones have poor expression of the cone marker PNA (Hasan *et al*., 2020), Fig. 1C). Immuno-histochemical analysis showed that subretinal injection of rAAV Gnat2::Lrit3 restored expression of LRIT3 as well as the cone marker stained by PNA to many cones (Fig. 1C). In addition to LRIT3 expression in cones (17 of 22 cone terminals in Fig. 1C, 3 marked by dotted circles) we also found a small number of rod terminals expressing LRIT3 (discrete puncta marked by arrows in Fig. 1iii). To investigate if the restored expression of LRIT3 rescued the missing signalplex components, mGluR6, GPR179, TRPM1, nyctalopin and RGS11, we immuno-stained retinal sections from control, *Lrit3*^*-/-*^, and rAAV Gnat2::Lrit3 injected *Lrit3*^*-/-*^ mice. Because there are no antibodies to nyctalopin, for its visualization we crossed a mouse transgenic line (Demas et al., 2006) that expresses a nyctalopin-EYFP fusion protein onto the *Lrit3*^*-/-*^ background and stained with antibodies to EYFP. Immunohistochemistry show that both nyctalopin-EYFP and TRPM1 are absent from *Lrit3*^*-/-*^ retinas, but expression is restored to the signalplex of DBCs connected to cones expressing LRIT3 (Fig 2. A,B) with patterns indistinguishable from controls. mGluR6 and GPR179 are both expressed normally in *Lrit3*^*-/-*^ rod BC terminals as evidenced by presence of puncta in *Lrit3*^*-/-*^ retinas (Fig. 3Aii, and Bii, respectively). In contrast, they are absent from the elongated profiles typical of the much larger cone terminals that have many invaginating ribbon synapses connecting to cone DBCs and flat synapses connecting to HBCs (Fig. 3). Following treatment with rAAV Gnat2::Lrit3, LRIT3 expression in the *Lrit3*^*-/-*^ retinas is present in many cones and mGluR6 and GPR179 expression also is restored to their postsynaptic DBC targets (Fig 3Aiii, Biii).

**Figure 2.**
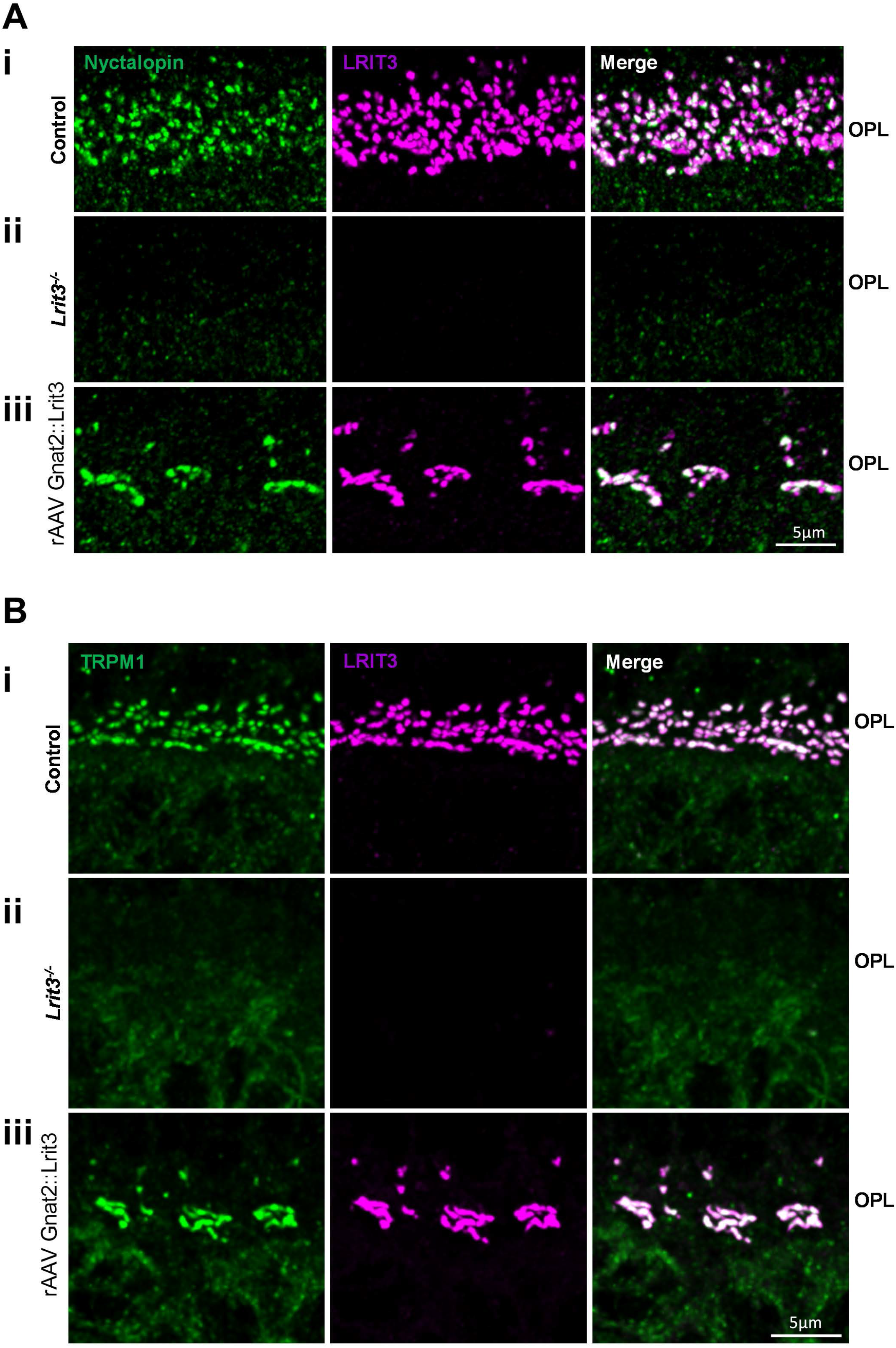
rAAV Gnat2::Lrit3 restores expression of Nyctalopin and TRPM1 to the dendrites of DBCs. (A) Transverse retinal sections of (Ai) control, (Aii) *Lrit3*^*-/-*^ and (Aiii) rAAV Gnat2::Lrit3 treated *Lrit3*^*-/-*^ retinas stained for Nyctalopin and LRIT3. Scale bar = 5µm. (B). Transverse retinal sections of (Bi) control, (Bii) *Lrit3*^*-/-*^ and (Biii) rAAV Gnat2::Lrit3 treated *Lrit3*^*-/-*^retinas stained for TRPM1 and LRIT3. Scale bar = 5µm.

**Figure 3.**
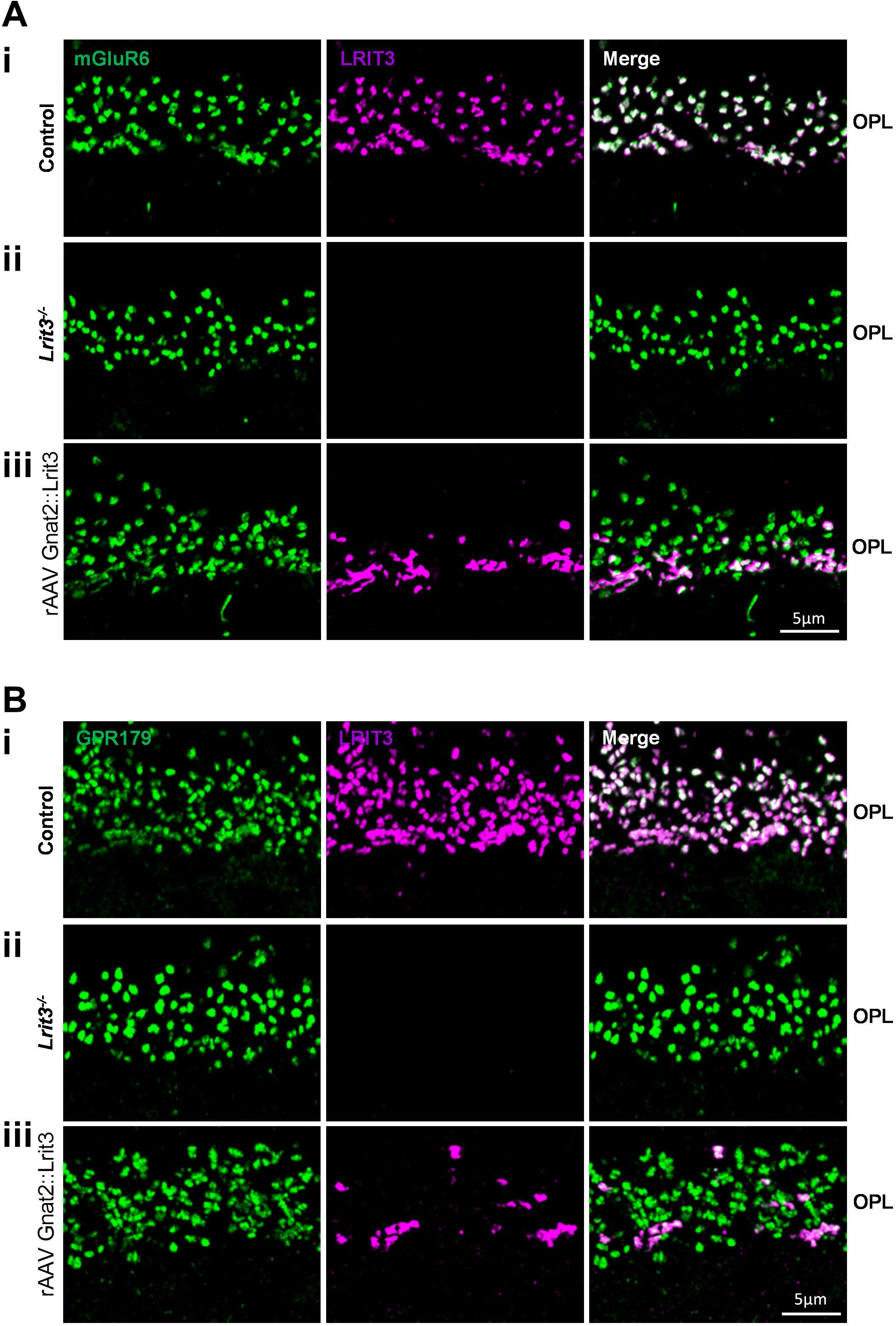
rAAV Gnat2::Lrit3 restores expression of mGluR6 and GPR179 to the dendrites of DBCs. (A) Transverse retinal sections of (Ai) control, (Aii) *Lrit3*^*-/-*^ and (Aiii) rAAV Gnat2::Lrit3 treated *Lrit3*^*-/-*^ retinas stained for mGluR6 and LRIT3. Scale bar = 5µm. (B). Transverse retinal sections of (Bi) control, (Bii) *Lrit3*^*-/-*^ and (Biii) rAAV Gnat2::Lrit3 treated *Lrit3*^*-/-*^ retinas stained for GPR179 and LRIT3. Scale bar = 5µm.

The small number of rods also expressing LRIT3 in the rAAV Gnat2::Lrit3 treated *Lrit3*^*-/-*^ retinas is a consistent finding, suggesting that the Gnat2 promoter used, while driving robust expression in cones, also has a low level of ectopic expression in a small number of rods, which leads to partial restoration of rod function (see Supplementary Fig. S3). We stained for RGS11 – one of two known RGS proteins utilized by the DBC signalplex, and found that it, too, is restored at the cone→cone DBC synapse (Supplementary Fig. S1). Finally, we stained for ELFN2, and as reported by others (Cao et al., 2020), found its expression is normal in the *Lrit3*^*-/-*^ mice (Supplementary Fig. S2). Combined, these data show that expression of LRIT3 in cones is necessary and sufficient to restore expression of nyctalopin, TRPM1, mGluR6, GPR179, and RGS11 to the cone DBC signalplex.

### rAAV-mediated LRIT3 expression in cones restores function to inner retina neurons in *Lrit3*^*-/-*^ mice

To assess restoration of cone DBC function after subretinal injection of rAAV Gnat2::Lrit3 in *Lrit3*^*-/-*^ retinas we used the photopic ERG. This initial assessment isolated the cone response by using a rod saturating background (20 cd s/m^2^). The photopic b-wave was measured using a 1.4 log cd s/m^2^ flash. In total we injected 32 *Lrit3*^*-/-*^ eyes (23 mice) with rAAV Gnat2::Lrit3 and 7 control eyes (4 mice) with rAAV CAG::GFP (Fig. 4A). The median response of rAAV Gnat2::Lrit3 injected eyes was 44µV, which is ∼33% of the rAAV CAG::GFP injected control eyes (median = 120µV, n=7 eyes, 4 mice; Fig. 4A). The characteristic lack of a b-wave in *Lrit3*^*-/-*^ mice is indicated by a single point in Fig. 4A for illustration purposes. Animals from this group then were randomized and used for further ERG, immunohistochemistry or electrophysiological assessment.

**Figure 4.**
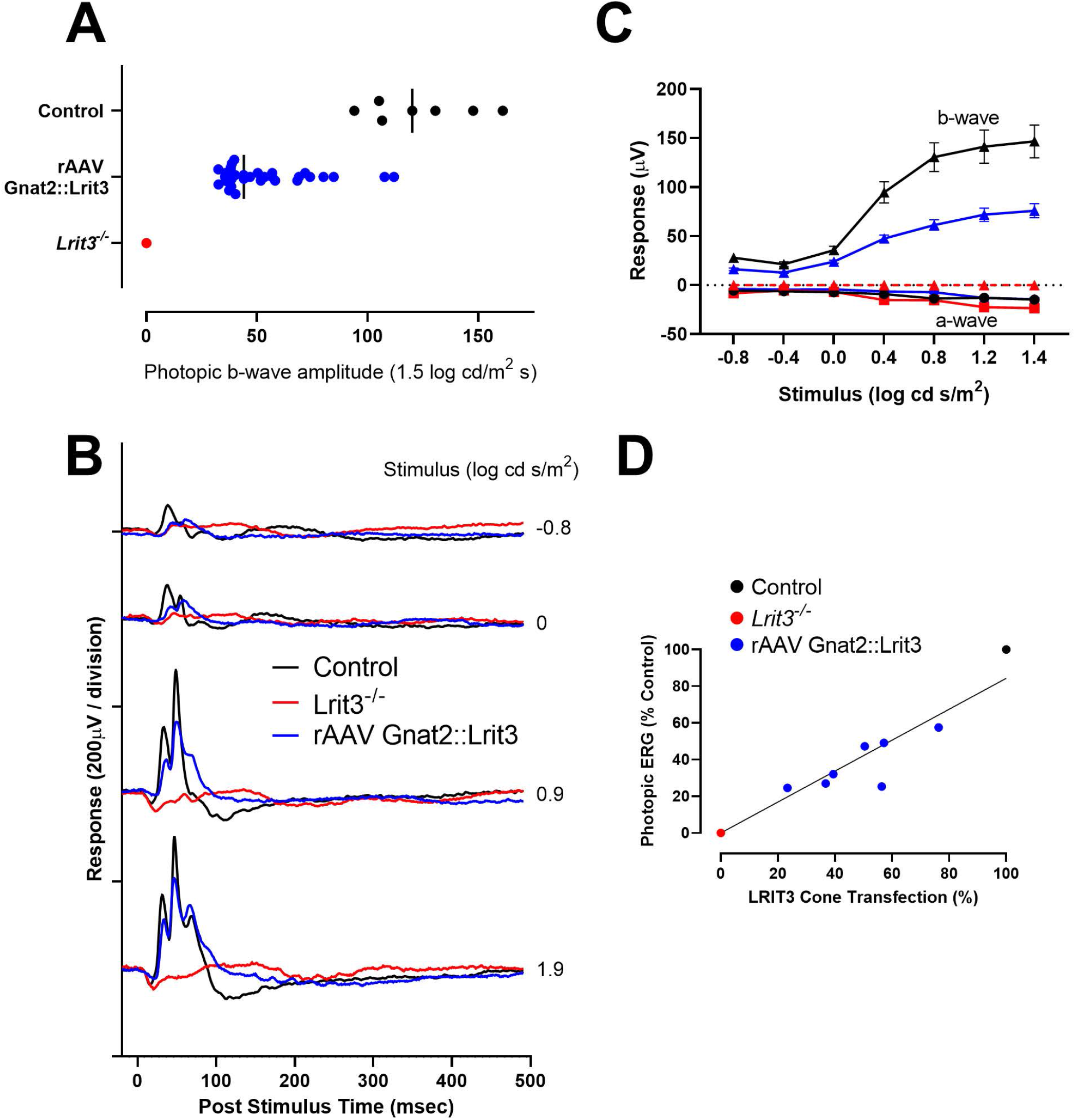
LRIT3 expression in *Lrit3*^*-/-*^ cones restores cone DBC function. (A) Photopic electroretinogram b-wave amplitude at 1.4 log cd s/m^2^ for control (7 eyes, 4 mice) and rAAV Gnat2::Lrit3 treated *Lrit3*^*-/-*^ eyes (32 eyes, 23 mice) mice. (B). Photopic electroretinograms at different stimulus intensities for one control eye (black), an age-matched *Lrit3*^*-/-*^ eye (red), and an rAAV Gnat2::Lrit3 treated *Lrit3*^*-/-*^ eye (blue). (C) The average stimulus-response plots for the ERG b-wave amplitudes under photopic conditions for control rAAV Gnat2::GFP (n=6, black), *Lrit3*^*-/-*^ (n=3, red) and rAAV Gnat2::Lrit3 treated *Lrit3*^*-/-*^ (n=8, blue). The dashed red lines indicate no b-wave (0 amplitude) in these animals. (D) Amplitude of the photopic b-wave as a function of the percentage of cones in *Lrit3*^*-/-*^ retinas infected with rAAV Gnat2::Lrit3. Control and *Lrit3*^*-/-*^ values are shown as a single point. Statistics: (C) 2-way repeated measures ANOVA; control b-wave responses were significantly greater than rAAV Gnat2::Lrtit3 treated for stimuli of 0.5 cd log cd s/m^2^ and above (p_adj_ <0.05), and rAAV Gnat2::Lrit3 treatment was significantly above baseline (that for *Lrit3*^*-/-*^) at 0.0 cd log cd s/m^2^ and above (p_adj_ < 0.01). ERG a-wave responses were not different between groups.

To assess if treatment also restored visual function, we obtained ERGs and measured amplitudes of the photopic ERG b-wave in control (rAAV CAG::GFP; n = 5) and treated (rAAV Gnat2::Lrit3; n = 6) *Lrit3*^*-/-*^ mice (Fig. 4). The photopic (Fig. 4B) and scotopic (Supplementary Fig. S3) waveforms for one control (rAAV CAG-GFP), one rAAV Gnat2::Lrit3 injected *Lrit3*^*-/-*^ and one uninjected *Lrit3*^*-/-*^ mouse, along with summary b-wave amplitude data from multiple animals (Fig. 4C) are shown. As previously reported (Neuille et al., 2014; Ray, 2013), *Lrit3*^*-/-*^ mice had neither a scotopic or photopic ERG b-wave. rAAV Gnat2::Lrit3 treated *Lrit3*^*-/-*^ mice, on the other hand, showed robust restoration of the cone DBC derived ERG waveform (Fig 4C) with near-complete recovery of the photopic b-wave at high intensities in some animals (example shown in Fig. 4B). The b-wave amplitude in rAAV Gnat2::Lrit3 eyes across the recorded population was always smaller than for control eyes (p_adj_<0.01 for all flash intensities, 2-way ANOVA, Tukey corrected). This result is consistent with the percentage of cones infected we observe (<100%, Fig. 4D). While the ERG response always exceeded the amplitudes observed in *Lrit3*^*-/-*^ eyes, the extent of cone infection and thus cone DBC rescue was variable, and this correlated with the amplitude of the photopic ERG b-wave (Fig. 4D, r^2^ = 0.87).

In addition to the restoration of the photopic (cone-driven) ERG b-wave we also found some recovery of the scotopic ERG b-wave (Supplementary Fig. S3). This result is consistent with our immunohistochemistry, which showed unambiguously that a small fraction of rods in the retina of treated rAAV Gnat2::Lrit3 treated *Lrit3*^*-/-*^ mice have restored rod BC signalplex proteins (Fig. 2,3), indicating low level ectopic expression of this vector /promoter combination in rod photoreceptors.

We conclude that infection with rAAV Gnat2::Lrit3 restores LRIT3 expression to infected cones as well as a small fraction of rods, and enables the functional assembly of signalplex proteins in connected cone and rod DBCs, rescuing their function as measured with full-field ERG.

### rAAV-mediated LRIT3 expression in cones restores light-evoked responses in *Lrit3*^*-/-*^RGCs

In the retina of control mice, short latency RGC responses (latency <100 msec) are evoked at light onset (ON) in ON-type ganglion cells, at light offset (OFF) in OFF-type ganglion cells, and at both light onset and offset in ON-OFF type ganglion cells. In all mouse models of cCSNB (*Grm6*^*-/-*^, *Trpm1*^*-/-*^, *Gpr179*^*nob5*^, *Nyx*^*nob*^, *Lrit3*^*nob6*^ and *Lrit3*^*-/-*^), all short-latency ON responses are absent (Demas *et al*., 2006; Neuille *et al*., 2017) (Hasan *et al*., 2020; Peachey et al., 2012; Renteria et al., 2006), consistent with the loss of light-evoked signaling in ON bipolar cells. Our previous work showed that OFF responses in *Lrit3*^*-/-*^ are also impacted and have smaller response amplitudes compared with control mice. The apparent causes are interference of OFF bipolar cell signaling from dysfunctional ON bipolar cells through cross-over inhibitory signaling pathways (Demb and Singer, 2015), and reduced excitatory drive from cones onto the OFF bipolar cells (Hasan *et al*., 2020).

The presence of b-wave oscillatory potentials in the example ERG traces (Fig. 4B, Supplementary Fig. S3) indicates restoration of visual function in the inner retina, both at photopic and scotopic light levels. To assess this at the level of the retinal output, we made targeted whole-cell electrophysiological recordings from identified ganglion cell types in retinas of control, *Lrit3*^*-/-*^, and rAAV Gnat2::Lrit3 treated *Lrit3*^*-/-*^ mice. We measured excitatory current responses in morphologically and functionally identified ON- and OFF-alpha RGCs and OFF- delta RGCs in retinal whole mounts under photopic (rod saturating) conditions. We postulated that the infection of cones by rAAV was highest near the injections site, so we recorded from ON-α RGCs located within a 300µm radius from the injection site while avoiding the injection scar (Fig. 5A).

**Figure 5.**
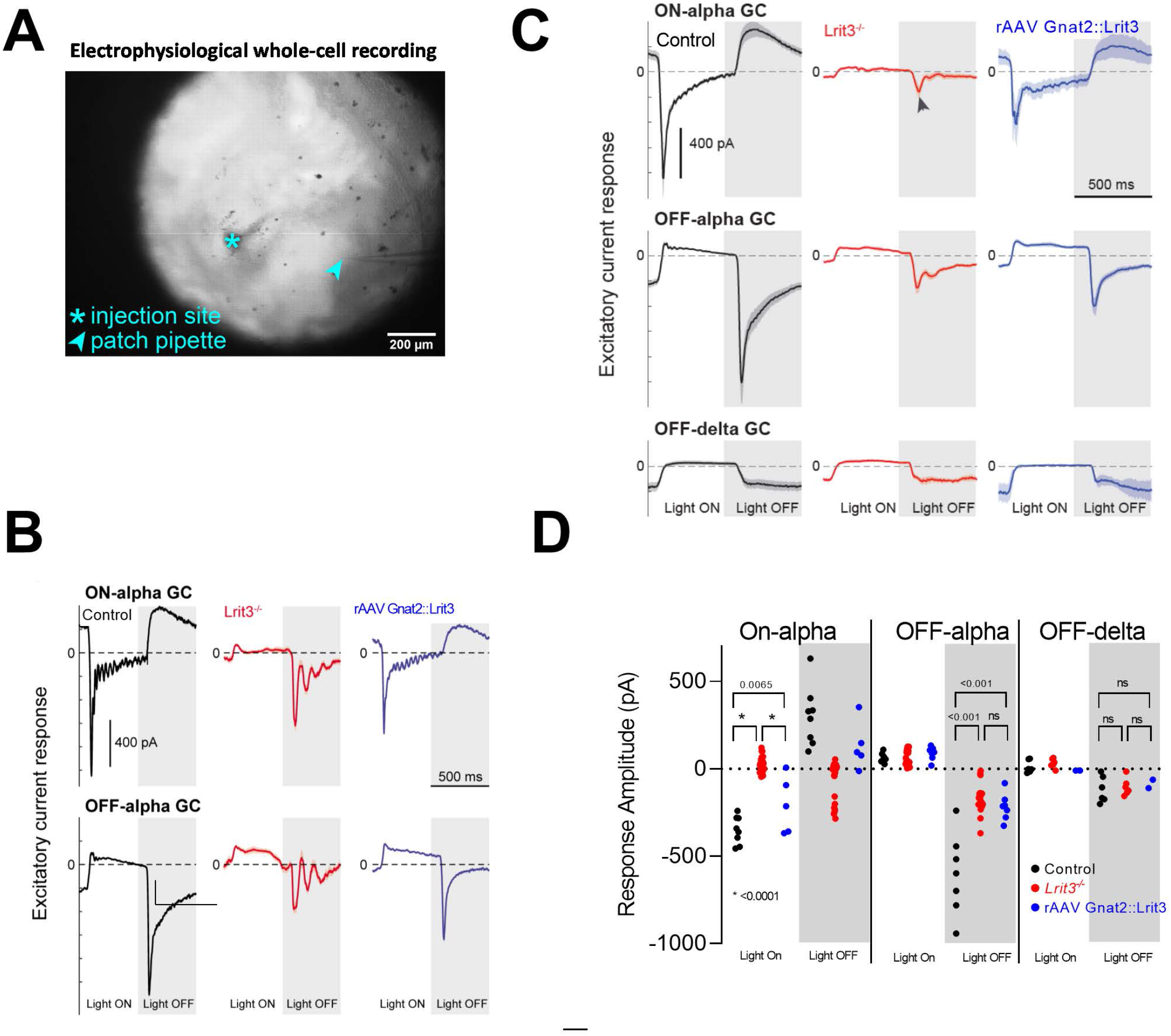
rAAV Gnat2::Lrit3 treatment restores visual function in ON alpha ganglion cells in *Lrit3*^*-/-*^ mice. (A) Brightfield microscope image of an ex vivo preparation of a rAAV Gnat2::Lrit3 treated *Lrit3*^*-/-*^ mouse retina. The tissue was mounted on opaque nitrocellulose filter paper with 1.2 mm diameter wholes under each retinal quadrant to enable brightfield imaging and visual stimulation of the photoreceptors through the microscope condenser light path (see Methods for details). The puncture site of the subretinal injection (asterisk) is readily visible and was avoided when targeting alpha-type ganglion cells for voltage clamp recording with a whole-cell recording pipette (arrowhead). (B) Single cell examples of excitatory currents (V_hold_ = -69 mV) recorded in ON (top) and OFF (bottom) alpha ganglion cells of control (black), *Lrit3*^*-/-*^ (red), and rAAV Gnat2::Lrit3 treated *Lrit3*^*-/-*^ mice (blue). The visual stimulus comprised a contrast reversing spot on a gray background (1Hz; 100% Michelson contrast). (C) As in B, traces show mean ± SEM for the recorded cell populations; sample sizes as indicated in D. (D) Response amplitudes of ON alpha, OFF alpha, and OFF delta ganglion cells following light increment of the contrast spot (light ON) and light decrement of the contrast spot (light OFF) in control (black), *Lrit3*^*-/-*^ (red), and rAAV Gnat2::Lrit3 treated *Lrit3*^*-/-*^ mice (blue).

Experiments in control ON-alpha GCs showed large amplitude (>0.4 nA), short latency excitatory currents at light onset, reflecting glutamatergic input from DBCs, and at light offset in an OFF-alpha GCs (representative example shown in Fig. 5B, population averages Fig. 5C; response amplitudes Fig. 5D). ON-α RGCs in *Lrit3*^*-/-*^ mice lacked a short latency ON response and instead showed an excitatory response and oscillations at light offset (Fig. 5B-D, (Hasan *et al*., 2020). OFF-alpha GCs in *Lrit3*^*-/-*^ had a reduced response amplitude and showed strong oscillations in the sustained excitatory light response (Fig. 5B-D). Responses in OFF-delta GCs were largely unaffected but showed some oscillations at light offset (Fig. 5C, D; (Hasan *et al*., 2020). rAAV Gnat2::Lrit3 treatment of *Lrit3*^*-/-*^ retinas restored short-latency excitatory responses at light onset in ON-alpha cells and eliminated the abnormal excitatory response at light offset (Fig. 5B-D). rAAV Gnat2::Lrit3 treatment further eliminated the abnormal oscillations at light offset in most OFF-alpha RGCs (Fig. 5B-D), however, the amplitude of the excitatory responses at light offset was still reduced, and similar to that of OFF-alpha RGCs in *Lrit3*^*-/-*^ retinas (Fig. 5D). Our small sample of OFF-delta RGCs in rAAV Gnat2::Lrit3 treated *Lrit3*^*-/-*^ retinas (n=2) were similar to those in control retinas (Fig. 5D, blue trace; n = 2). We conclude that restoration of LRIT3 expression at the cone→cone DBC synapse in *Lrit3*^*-/-*^ mice reverses the impaired light signaling in the ON-alpha type retinal ganglion cells assayed here.

Although LRIT3 plays a major role in signalplex assembly and function of the ON signaling pathway, our previous study showed that signaling in the retinal OFF pathway in *Lrit3*^*-/-*^ mice was also impaired (Hasan *et al*., 2020). Specifically, the response amplitude of Type 1 OFF BCs (BC1s) were also abnormal. To determine if rAAV Gnat2::Lrit3 treatment of *Lrit3*^*-/-*^ retinas restored normal function in OFF BCs, we made targeted whole-cell recordings of genetically identified BC1s in control, *Lrit3*^*-/-*^, and rAAV Gnat2::Lrit3 treated *Lrit3*^*-/-*^ retinas.

BC1 recordings from control and *Lrit3*^*-/-*^ retinas recapitulated our published data (Fig. 6A, B). Control BC1s showed ∼50 pA amplitude excitatory current at light off and suppression of this excitation at light on; ∼30pA amplitude release of inhibition at light OFF and ∼60 pA inhibition at light on (n = 3). *Lrit3*^*-/-*^ BC1s showed no significant change in excitation at light off or light on, spontaneous inhibition consistent with oscillations at light off, and suppression of inhibition at light on (Fig. 6A, B). rAAV Gnat2::Lrit3 treated *Lrit3*^*-/-*^ retinas showed restoration of responses in some, but not in all recorded BC1s (Fig. 6D1, D2). This variability may be explained by regional differences in efficacy of cone rAAV infection, which could not be visualized in the electrophysiological experiments. Importantly, we found that about half of the sampled BC1s had substantial (>20 pA) light evoked responses with limited or no oscillations in the excitatory or inhibitory current (Fig. 6 D).

**Figure 6.**
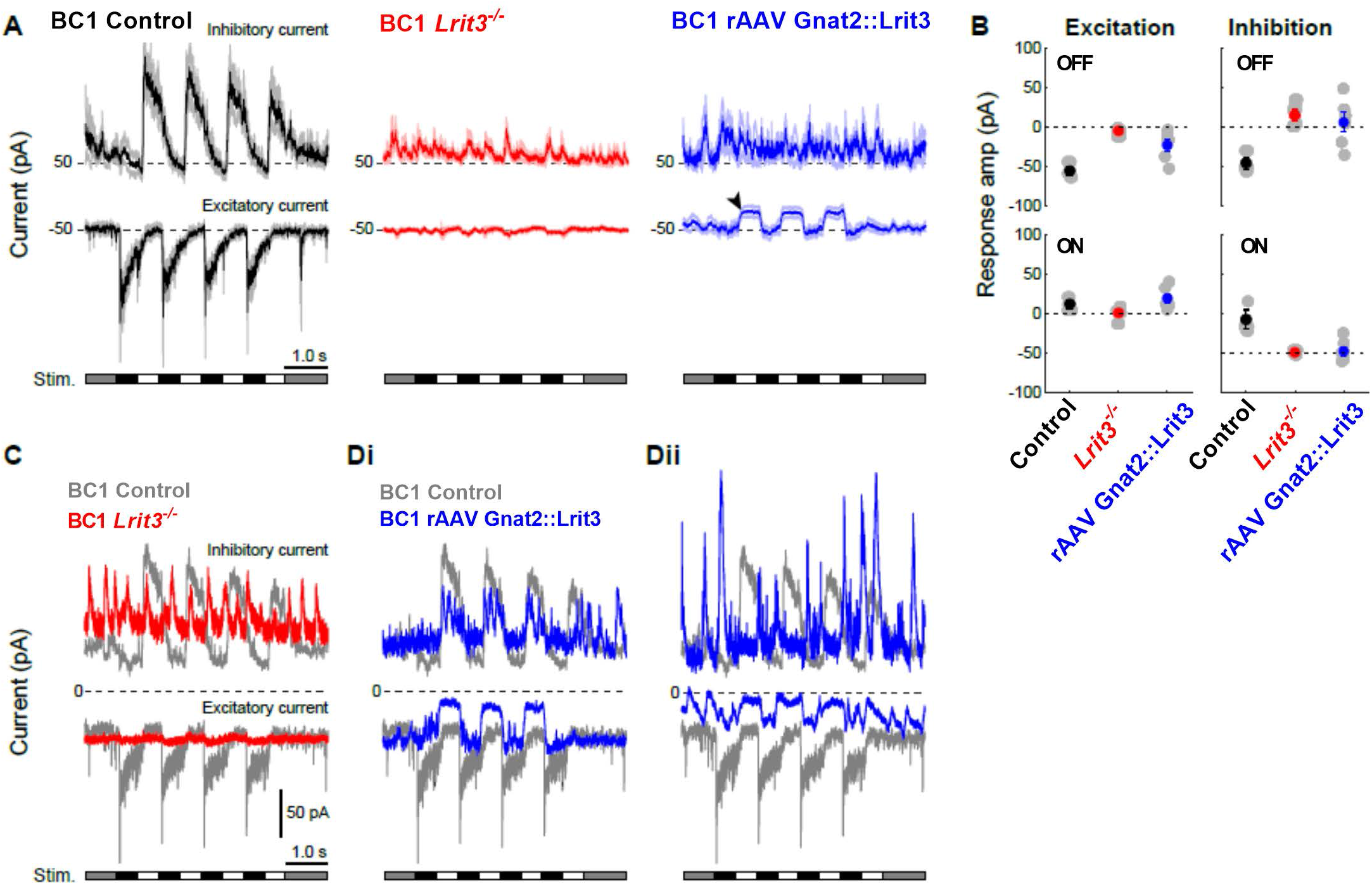
rAAV Gnat2::Lrit3 treatment restores visual function in some OFF bipolar cells in *Lrit3*^*-/-*^ mice. (A) Population average responses of OFF type (BC1) bipolar cells obtained with targeted whole-cell electrophysiological recordings. Traces show inhibitory synaptic current (top, V_hold_ = 0 mV) and excitatory synaptic current (bottom; V_hold_ = -69 mV) recorded during visual stimulation with a contrast reversing spot on a gray background (100% Michelson contrast; 150 µm diameter; 1 Hz; time course indicated below). Control, n = 3; *Lrit3*^*-/-*^, n = 6; rAAV Gnat2::Lrit3 *Lrit3*^*-/-*^ treated, n = 6. (B) Current amplitudes of the excitatory (left) and inhibitory (right) response phases following the switch to dark (OFF, top) and light (ON, bottom) spot stimulus period. (C) Single cell example of a BC1 in *Lrit3*^*-/-*^ retina showing negligible stimulus-evoked excitation (bottom, red trace) compared with a BC1 in control, (bottom, gray trace) and oscillatory inhibitory currents with no apparent stimulus-evoked modulation (red top trace). (D) rAAV Gnat2::Lrit3 mediated LRIT3 expression in *Lrit3*^*-/-*^ mice caused restoration of stimulus-evoked responses in excitatory (top, blue traces) and inhibitory currents (bottom, blue traces) and loss of oscillating inhibitory input to varying degree. Panels show single cell examples of a BC1 with stronger rescue (Di) and weaker rescue (Dii), representative of the observed range.

Together, these data show that rAAV Gnat2::Lrit3 mediated expression of LRIT3 in *Lrit3*^*-/-*^ cones can restore signaling in cone DBCs, OFF BCs and downstream ON and OFF RGCs.

## DISCUSSION

The need for LRIT3 in organizing the DBC signalplex differs between rod and cone pathways. Our results demonstrate for the first time that rAAV mediated restoration of expression of LRIT3 in cone photoreceptors trans-synaptically organizes the post-synaptic signaling complex in cone DBCs that mediates excitatory inputs to downstream circuitry.

At the cone to cone DBC synapse, LRIT3 is required for localization of mGluR6, GPR179, TRPM1 and Nyctalopin (Hasan *et al*., 2019; Hasan *et al*., 2020). In contrast, at the rod to rod BC synapse, LRIT3 is required only for the localization of the signalplex proteins nyctalopin and TRPM1. Therefore, we postulated that this difference may be due to expression of LRIT3 in both in cones and in cone BCs. However, our results are not consistent with this hypothesis, as we demonstrate that presynaptic expression in cones is sufficient for assembly of the cone DBC signalplex. This is demonstrated by the return of both signalplex expression and functional rescue of the ERG b-wave and visually evoked responses in ON RGCs.

At the cone to cone DBC synapse a second LRR containing protein, ELFN2, or ELFN1 in ELFN2 knockout mice, also is required for assembly of mGluR6 and GPR179 into the signalplex (Cao *et al*., 2020). The dependency of TRPM1 on ELFN2(ELFN1) is currently unknown. ELFN2 is expressed normally in *Lri3*^*-/-*^ mice (this study, (Cao *et al*., 2020) and LRIT3 is expressed normally in ELFN2/1 double KO mice (Cao *et al*., 2020). Thus, these data imply that both LRIT3 and ELFN2 (or ELFN1) are required for normal cone DBC signalplex assembly and function, and that they likely represent members of a presynaptic complex that trans-synaptically organizes post-synaptic elements to maximize function. The difference in the impact of the loss of LRIT3 on rod DBCs and Cone DBCs also suggests that a different pre-synaptic complex is required to assemble the rod BC signalplex.

The loss of LRIT3 from rods results in only loss of TRPM1 and nyctalopin from the rod BC signalplex, meaning that remaining proteins are capable of localizing mGluR6 and GPR179. Two candidates are LRRTM4 (Agosto and Wensel, 2021) and ELFN1 (Cao et al., 2015), both of which colocalize with the rod DBC signalplex. LRRTM4 is a LRR containing protein whose removal in mice does not impact rod BC function (Sinha et al., 2020). Whether other family members compensate for this loss in the knockouts is unknown. The situation at the rod to rod BC synapse for ELFN1 is complex. Constitutive knockout of ELFN1 disrupts the structure of the rod spherule during development and triggers dendritic extensions of rod BC and HC dendrites into the ONL (Cao *et al*., 2015). ELFN1 also interacts with the α2δ4 subunit of the rod voltage gated calcium channels critical for glutamate release (Wang et al., 2017). The phenotype of the ELFN1 knockout is therefore consistent with other mutations impacting the channels function, CACN1F (α1F), CACNA2D4 (α2δ4), CACNB2 (β2), CABP4 as well as bassoon and ribeye (Ball et al., 2002; Dick et al., 2003; Haeseleer et al., 2004; Kerov et al., 2018; Mansergh et al., 2005; Maxeiner et al., 2016; Wang *et al*., 2017). In the mature retina ELFN1 does not co-localize with α2δ4, but when removed from adult retina using a tamoxifen conditional knockout strategy, mGluR6 also is lost (Cao *et al*., 2020), suggesting ELFN1 may be required for the normal localization of mGluR6, separate from its role during development. Together, these data suggest that LRIT3 is required for TRPM1/Nyctalopin colocalization on rod BCs, and that ELFN1 is critical for localization of mGluR6. Based on these data we postulate that both are required as part of a presynaptic complex that correctly assembles the post-synaptic rod BC signalplex.

Retinas in *Lrit3*^*-/-*^ lack signally to DBCs (Hasan *et al*., 2018; Hasan *et al*., 2020; Neuille *et al*., 2015; Ray, 2013) and as a consequence inputs via the ON pathway (Hasan *et al*., 2019; Hasan *et al*., 2020). A second consequence of the loss of LRIT3 and ON pathway signaling is that RGCs exhibit spontaneous oscillating activity (Hasan *et al*., 2020), which are evident in the patch clamp recordings (Fig 5). This is consistent with reports that RGCs in two other cCSNB models, *Nyx*^*nob*^ and *Trpm1*^*-/-*^, that cause loss of TRPM1 also exhibit this oscillating activity (Demas *et al*., 2006; Takeuchi et al., 2018; Winkelman et al., 2019). Oscillatory activity also has been reported in mouse models of retinitis pigmentosa, which lose photoreceptor inputs to DBCs, although the oscillation frequency is different than in the cCSNB models, (Borowska et al., 2011; Stasheff, 2008; Trenholm et al., 2012). Given this abnormal retinal activity and the fact it could be associated with other clinical complications in CSNB patients, such as nystagmus, we asked if correcting the phenotype in some cones restores function and eliminates the oscillations. We show that this is indeed the case, as oscillations are absent from the ON alpha RGCs in rAAV Gnat2::Lrit3 treated *Lrit3*^*-/-*^ mouse retinas (Fig. 5). Restoring function to rod BCs in *Lrit3*^*-/-*^ mice also eliminates the oscillations in RGCs *Lrit3*^*-/-*^ mice (Hasan *et al*., 2020). Thus, given restoring function in either rod BCs or cone DBC eliminates these oscillations, this is consistent with the hypothesis that the AII amacrine cell is involved in the abnormal oscillatory activity (Borowska *et al*., 2011; Trenholm *et al*., 2012).

Loss of LRIT3 in mice, humans, and dogs causes cCSNB (Neuille *et al*., 2014; Zeitz et al., 2013) (Das et al., 2019; Ray, 2013). cCSNB in mice and humans is also caused by loss of other signalplex proteins, including nyctalopin, mGluR6, and GPR179. A characteristic of this disorder in mouse models, and presumably humans, is that the rod and cone synaptic architecture is normal. cCSNB, therefore, provides an ideal situation in which to restore function using gene therapy approaches. In this and our previous studies (Hasan *et al*., 2019; Hasan *et al*., 2020) we show that expression of LRIT3 in photoreceptors of adult *Lrit3*^*-/-*^ mice restored retinal function. The degree of rescue depends on the number of photoreceptors that can be transduced, which in the current study ranged from 23-76% of the cones, although the coverage was not uniform. This is highlighted by the fact that when analyzing BCs and RGC function the degree of rescue was variable between cells (Fig 5,6), presumably caused by differences in the fraction of restored cones/cone DBCs providing input to an individual recorded RGC.

The functional rescue of the CSNB phenotype by expressing LRIT3 in rods and cones supports a role of LRIT3 as a trans-synaptic organizer of the DBC signalplex, although two reports suggest partial rescue may be possible by expressing LRIT3 in DBCs. Zeitz and colleagues used AAV mediated expression of LRIT3 using DBC or photoreceptor specific promoters to restore function in *Lrit3*^*nob6*^ mice (Varin *et al*., 2021). Some ERG rescue was present in animals when LRIT3 expression was targeted to DBCs, although the extent of scotopic ERG rescue was greater when photoreceptors were targeted (Varin *et al*., 2021). In another study LRIT3 was targeted to DBCs in a *Lrit3*^*-/-*^ canine model of cCSNB (Miyadera *et al*., 2022. This partially restored the scotopic ERG b-wave, and resulted in recovery of visual function under scotopic conditions as assessed by behavioral tests (Das *et al*., 2019; Miyadera *et al*., 2022). In neither of these studies was there any rescue of cone DBC function as assessed by photopic ERGs. While the reason remains unclear, one possible explanation is poor transduction of cone DBCs by the AAV2.7M8 and AAV2^K9#4^ vectors used in the mouse and canine models, respectively. Another possibility is that there is low-level ectopic expression of the vectors in rods, which are 20 fold more numerous than cones, and at least 10 fold more numerous than DBCs, which could restore scotopic function. As we have seen in this study restoring function in small numbers of rods can restore the ERG b-wave in *Lrit3*^*-/-*^ mice. Resolution of this pre-versus post-synaptic expression awaits further studies in which imaging resolution is sufficient to establish pre- or post-synaptic expression, a question that remains for future studies.

The current study is the first to show that expression of LRIT3 in cones can restore all the key elements (nyctalopin, mGluR6, GPR179, and TRPM1) required for signal transduction in cone DBCs and their function as measured by the ERG and light adapted recordings of RGCs. This adds to a growing body of research establishing that there is a complex interacting network of LRR-containing proteins at photoreceptor synapses with DBCs that assemble the postsynaptic excitatory receptor complex (see (Furukawa *et al*., 2020) for review). These now include Nyctalopin, LRIT3, ELFN1 and ELFN2, LRIT1 and Latrophilin. It is becoming increasingly clear that the exact makeup of this network is different at rod compared with cone synapses.

LRR containing proteins such as LRIT3 are part of a large family of proteins that includes NGLs, LRRTMs, Slitrks, SALMS, and FLRTs – many of which are known to be involved in trans-synaptic organization throughout the CNS (see review (Schroeder and de Wit, 2018)). While their exact modes of action remain poorly understood, these individual proteins have been shown to be involved in regulation of synaptic strength, spine density, synapse density, and synaptic plasticity. Perhaps not surprisingly, many have been associated with disorders such as autism, schizophrenia, and Alzheimer’s Disease (see review by Schoeder and de Wit, 2018). The results reported here make it clear that LRR containing proteins are critical to assembling the post-synaptic metabotropic glutamate receptor 6 complex for visual function of DBCs in the retina. This renders these results highly relevant to organization of similar signaling systems present throughout the rest of the brain, particularly given the known role of synaptic dysfunction in human neurological diseases.

## Acknowledgements

This work was supported by funding from the NIH (R01 EY12354 to R.G.G. and N.H., and R01EY028188 to B.G.B), and support from the Preston Pope Joyes Endowed Chair in Biochemical Research (R.G.G.). Technical assistance was provided by Timothy Hoffman and Kynan Jarret.

## Author Contributions

Conceptualization, R.G.G., N.H. and B.G.B; Investigation, N.H., B.G.B Writing – Original Draft, R.G.G.; Writing – Review and Editing, R.G.G., N.H, B.G.B.; Funding Acquisition, R.G.G., N.H. and B.G.B.; Supervision, R.G.G., B.G.B.

## STAR METHODS

### Animals

All procedures were performed in accordance with the Society for Neuroscience policies on the use of animals in research and the University of Louisville Institutional Animal Care and Use Committee. Animals were housed in the University of Louisville AALAC approved facility under a 12 h/12 h light/dark cycle. The phenotypes of all the mouse lines have been previously published. *Lrit3*^*emrgg1*^, is referred to as *Lrit3*^*-/-*^ (Hasan *et al*., 2019) *Trpm1*^*-/-*^, *(Trpm1*^*tm1Lex*^), (Shen *et al*., 2009); *Grm6*^*-/-*^ (Masu et al., 1995); *Nyx*^*nob*^ (Gregg et al., 2003); *GPR179*^*nob5*^ (Peachey *et al*., 2012); *TgEYFP-Nyx* (Gregg et al., 2005); *MitoP-CFP* (Misgeld et al., 2007).

*Lrit3*^*-/-*^ and C57Bl/6J as controls, of both sexes, were used throughout. For all procedures, mice were anesthetized with a ketamine/xylazine solution (118/11 mg/kg, respectively) diluted in normal mouse Ringer’s prior to subretinal injections and ERG recordings. Mice were euthanized by CO_2_ exposure according to AVMA guidelines. Data from mice in which there was gross retinal damage from the injection procedure were excluded from the analyses.

### Antibodies

Antibodies used to label signalplex proteins have been described previously (Hasan *et al*., 2019) and are listed in Supplementary Table 1. Specificity of all antibodies was validated on sections from the respective knockout mouse retinas.

### Retina preparation for immunohistochemistry

Retina sections and wholemounts were prepared as described previously (Hasan *et al*., 2019). Briefly, mice were killed by CO_2_ inhalation followed by cervical dislocation, the eyes were enucleated and the cornea and lens removed. Retinas were dissected in PBS (pH 7.4) and fixed for 15-30 mins in phosphate buffer (0.1M PB) containing 4% paraformaldehyde, washed in PBS (3x 10 mins), and cryoprotected in a graded series of sucrose solutions (5, 10, 15 and 20% in 0.1M PB) and finally in OCT:20% sucrose (2:1). Cryoprotected retinas were frozen, and transverse 18μm sections were cut on a cryostat (Leica Biosystems, Buffalo Grove, IL) and mounted on Superfrost Plus slides (Thermo Fisher Scientific, Waltham, MA), and stored at - 80°C.

Immunohistochemistry methods have been described previously (Hasan et al., 2016). Sections were imaged on an FV-3000 Confocal Microscope (Olympus) and contrast and brightness adjusted using Fluoview Software (Olympus, Waltham, MA) or Photoshop (Adobe Systems, San Jose, CA).

### rAAV Production and Injection

To express LRIT3 in cones, we used rAAVs with gene expression controlled by the ProA1 promoter, which is part of the *Gnat2* promoter (Drinnenberg *et al*., 2018). Hereafter we refer to this promoter as Gnat2. The Lrit3 expression construct was packaged in the rAAV8 capsid by Vigene Biosciences (now Charles River). The virus expressing LRIT3 under the control of the Gnat2 promoter is referred to as rAAV Gnat2::Lrit3. rAAVs were introduced into the retinas of postnatal day 35-40 (P35-P40) mice by subretinal injection (1.0µl of 1 × 10^13^ vg/ml) using a specialized syringe (www.borghuisinstruments.com). Retinal function and protein expression were assessed four to eight weeks after rAAV injection.

### Electroretinography

Electroretinogram (ERG) methods were as described previously (Ray et al., 2014). Mice were dark adapted overnight, anesthetized with a ketamine/xylazine solution (118/11 mg/kg, respectively), and prepared for ERG recordings under dim red light. Pupils were dilated and accommodation relaxed with topical applications of 0.625% phenylephrine hydrochloride and 0.25% Tropicamide and the corneal surface anesthetized using 1% proparacaine HCl. Body temperature was maintained using a feedback-controlled electric heating pad (TC1000; CWE Inc.). A contact lens with a gold electrode (LKC Technologies Inc.) was placed on the cornea and wet with artificial tears (Tears Again; OCuSOFT, Gaithersburg, MD). Ground and reference needle electrodes were placed in the tail and on the midline of the forehead, respectively. Scotopic responses were measured with test flashes (from -3.6 to 1.8 log cd s/m^2^) presented to dark adapted animals; photopic responses were measured with test flashes (from -0.8 to 1.9 log cd s/m2) after 5 min light adaption to a rod-saturating background (20 cd/m^2^).

### RGC and OFF BC whole-cell recordings from retinal whole-mounts

Whole-cell voltage clamp recordings of BC1 and ON- and OFF alpha and OFF delta RGCs were obtained as described previously (Borghuis et al., 2014). BC1 cells were recorded with cesium-based internal pipette solution in the whole mount retina of MitoP-CFP transgenic mice on either a *Lrit3*^*-/-*^ or control C57Bl/6 genetic background. RGCs were targeted for recording based on soma size in *Lrit3*^*-/-*^ and control retinas. Cell type was verified functionally by signature features of the light-evoked current responses during recording, and confirmed anatomically post-hoc using two-photon fluorescence imaging of dye fills (Sulphorhodamine 101). The retina was stimulated at photopic light level (∼1.0 × 10^4^ photoisomerizations (R*) cone^-1^ sec^-1^) using a modified LED video p _max_ = 395 nm; HP). Light-evoked excitatory and inhibitory currents were recorded in whole-cell configuration at the reversal potential for chloride (−69 mV) and cations (0 mV), respectively, using conventional methods (Hasan *et al*., 2020).

### Statistical analyses

Prism 9.4.1 (681) (Graphpad Software, Inc., La Jolla, CA) was used to perform the appropriate statistical analyses as indicated in the text and figure legends. Tukey post-hoc tests were used to adjust for multiple testing when appropriate and the p_adj_ is reported. Statistical significance, p≤ 0.05.

---

**Supplementary Table 1.**
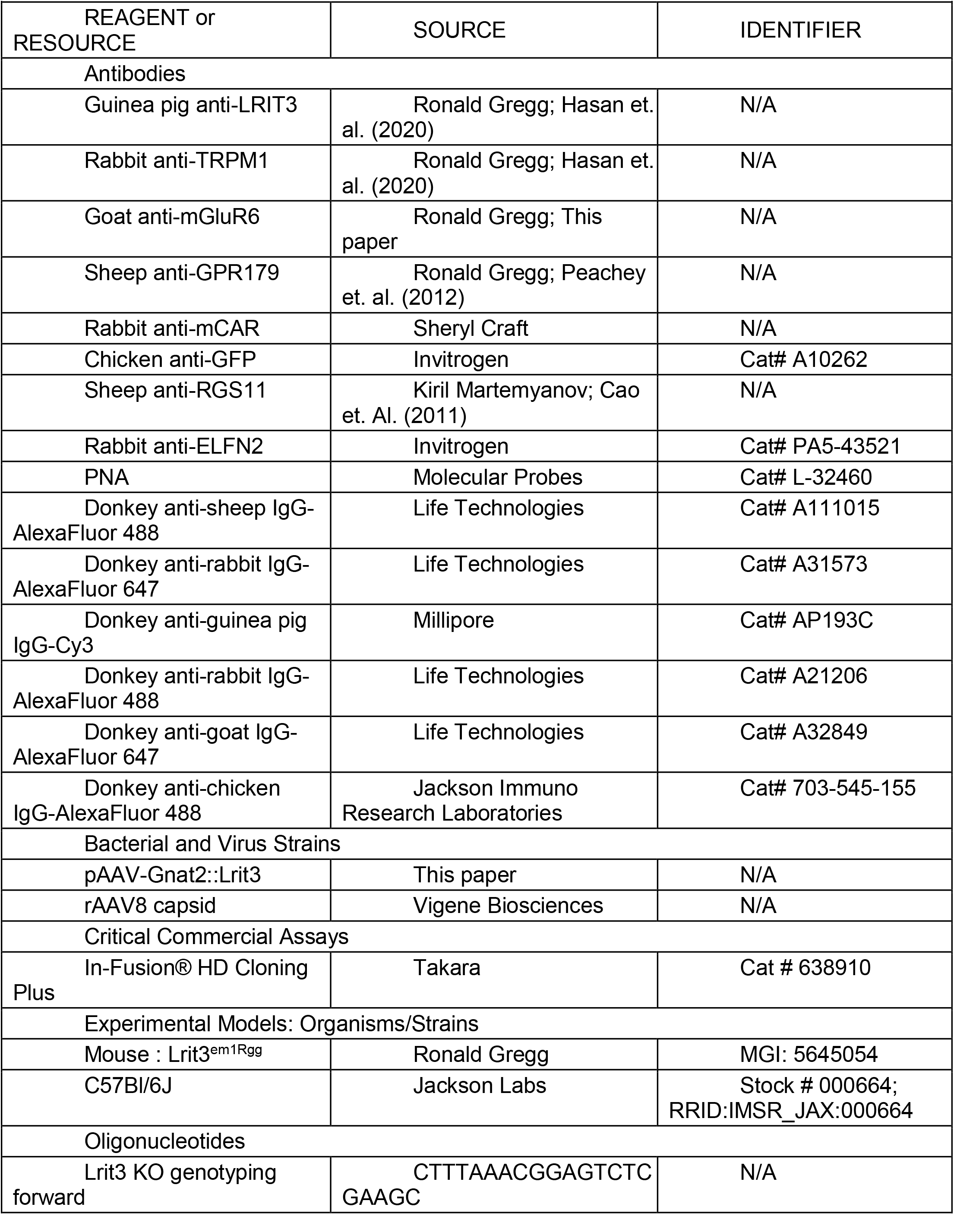
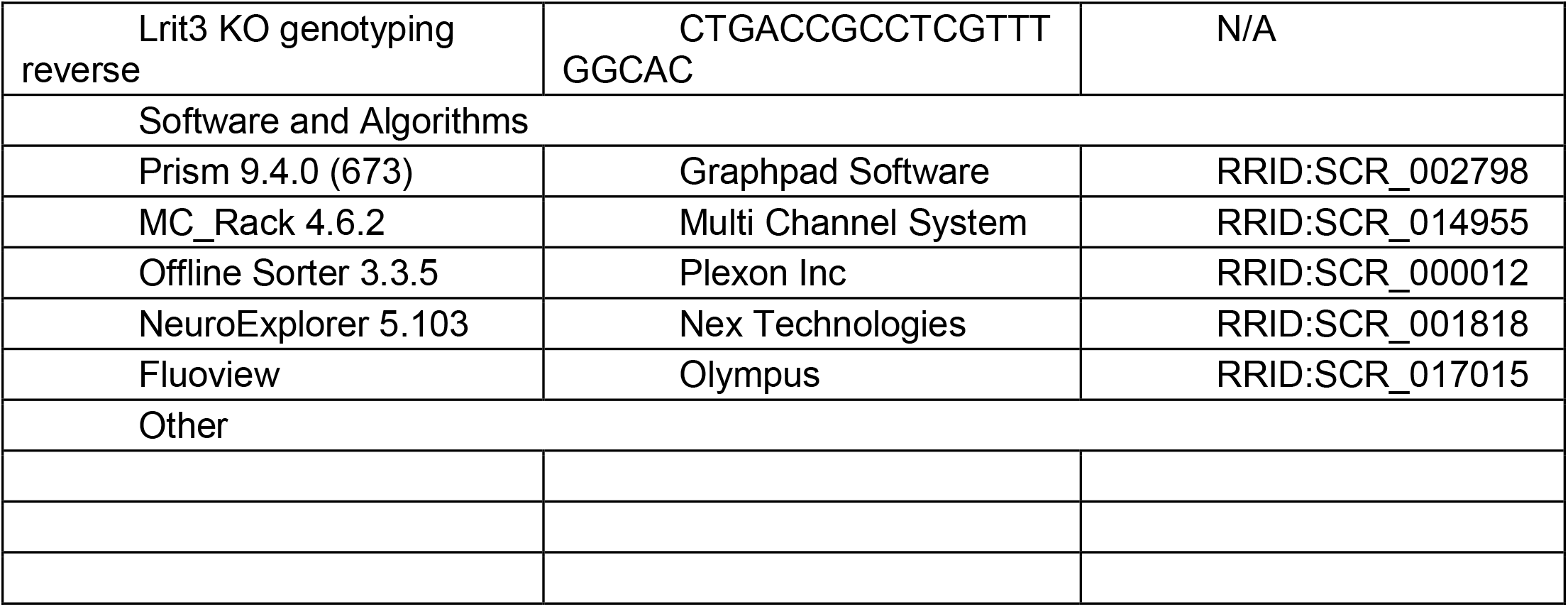
Key Resources.

