## Supplementary Figures for "LRIT3 expression in cone photoreceptors restores post-synaptic bipolar cell signalplex assembly and function in *Lrit3^-/-^* mice"

### Supplementary Fig. S1

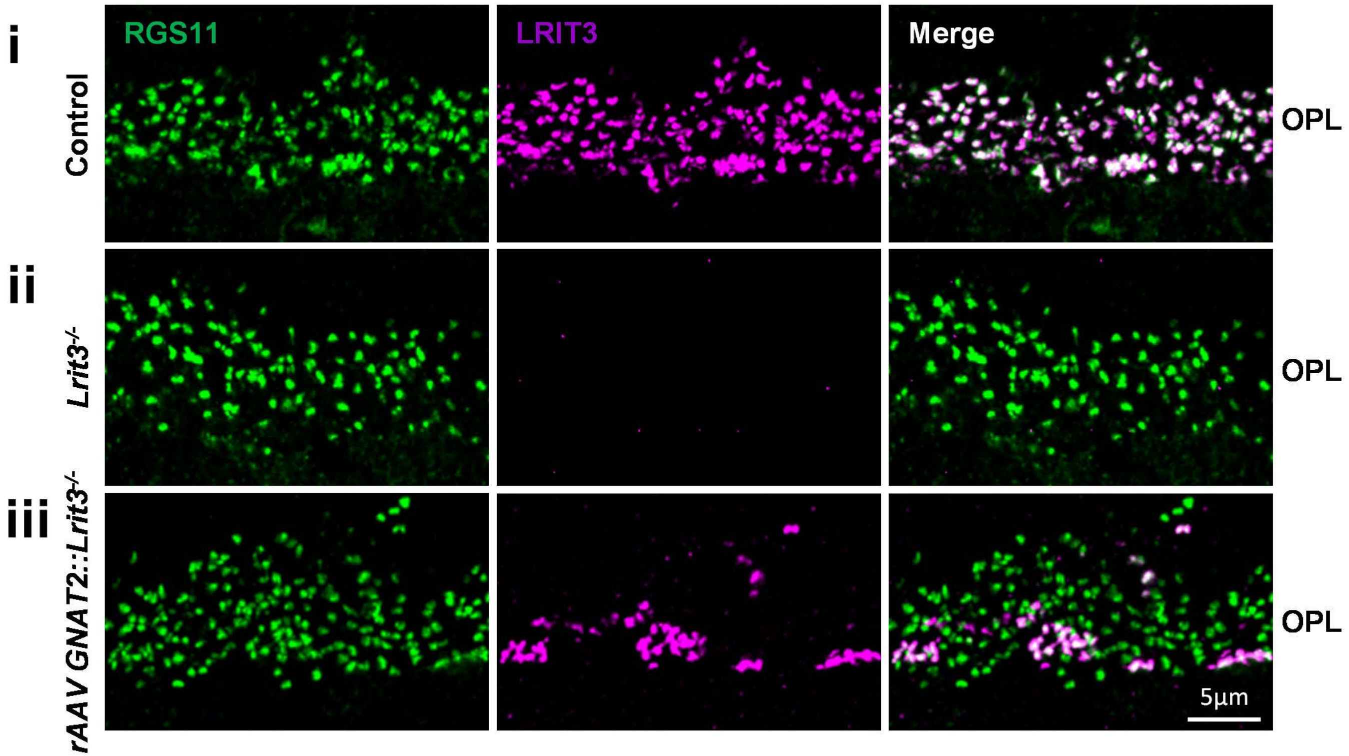


**Supplementary Figure S1. RGS11 expression is restored in cone DBCs connected to LRIT3 expressingn cones.** Transverse retinal sections of (i) control, (ii) *Lrit3*$^{-/-}$ and (iii) rAAV Gnat2::Lrit3 treated *Lrit3*$^{-/-}$ retinas stained for LRIT3 and RGS11. Scale bar = 5µm.

### Supplementary Fig. S2

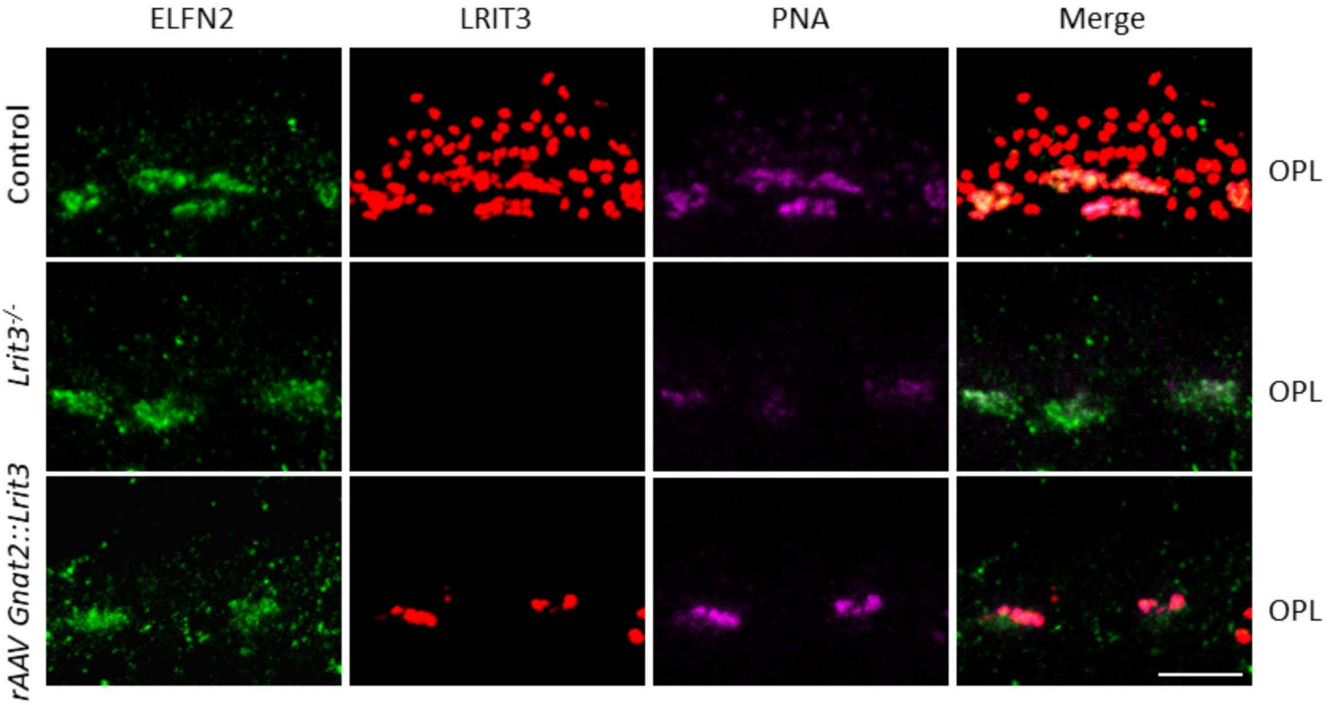


**Supplementary Figure S2. ELFN2 expression is retained in *Lrit3*$^{-/-}$ retinas.**
Transverse retinal sections of (i) control, (ii) *Lrit3*$^{-/-}$ and (iii) rAAV Gnat2::Lrit3 treated *Lrit3*$^{-/-}$ retinas stained for ELFN2, LRIT3, PNA, and the merged image. Scale bar = 5µm.

### Supplementary Fig. S3

#### Scotopic ERG

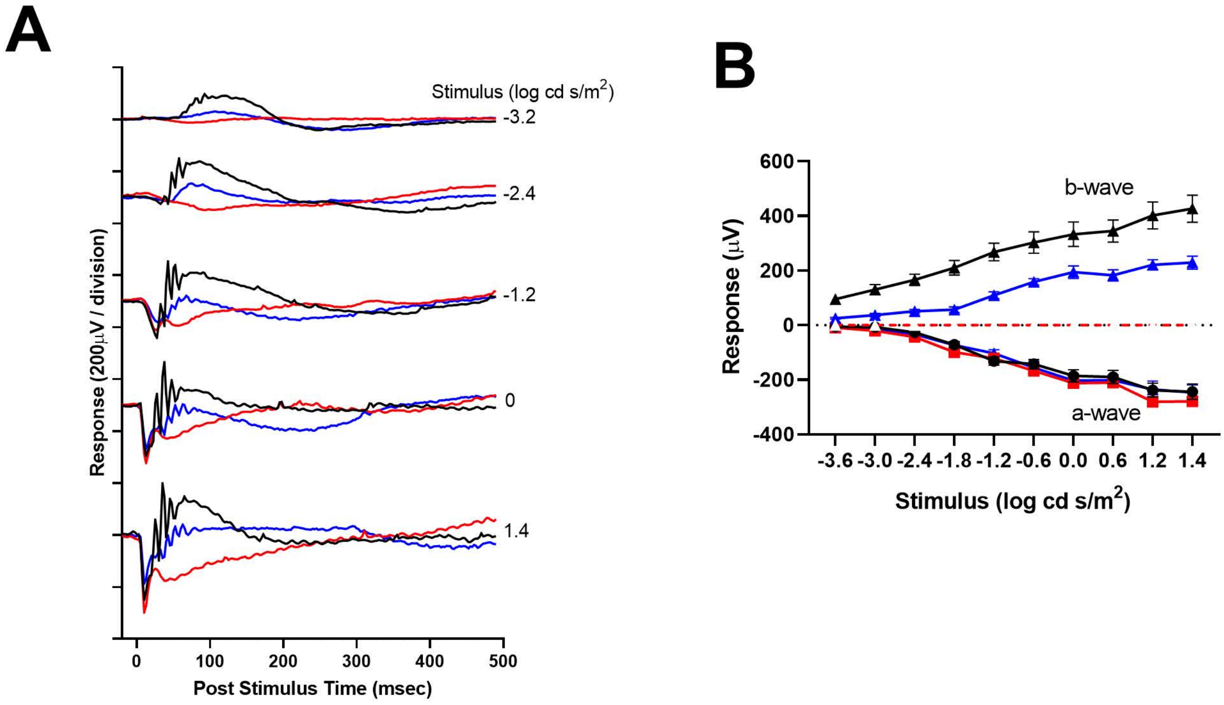


**Supplementary Fig S3. Low level LRIT3 expression in *Lrit3*$^{-/-}$ rods restores rod BC function**. (A) Scotopic electroretinograms at different stimulus intensities for one control eye (black), an age-matched *Lrit3*$^{-/-}$ eye (red), and an rAAV Gnat2::Lrit3 treated *Lrit3*$^{-/-}$ eye (blue). (B) The average stimulus-response plots for the ERG a- and b-wave amplitudes under scotopic conditions for control rAAV Gnat2::GFP (n=6, black), *Lrit3*$^{-/-}$ (n=3, red) and rAAV Gnat2::Lrit3 treated *Lrit3*$^{-/-}$ (n=8, blue). The dashed red lines indicate no b-wave (0 amplitude) Statistics: 2-way repeated measures ANOVA; control responses were significantly greater than rAAV Gnat2::Lrtit3 treatment at all flash intensities ($p_{adj} < 0.004$), responses in rAAV Gnat2::Lrit3 treated *Lrit3*$^{-/-}$ were significantly above baseline (that for *Lrit3*$^{-/-}$) at and above -1.8 cd log cd s/m2 ($p_{adj} < 0.0004$); a-wave responses were not different between groups.
